# Mixed Logistic Regression in Genome-Wide Association Studies

**DOI:** 10.1101/2020.01.17.910109

**Authors:** Jacqueline Milet, Hervé Perdry

**Affiliations:** Université de Paris, MERIT, IRD, Paris, F-75006, France; CESP Inserm, U1018, UFR Médecine, Univ Paris-Sud, Université Paris-Saclay, Villejuif, France

## Abstract

**Motivation:** Mixed linear models (MLM) have been widely used to account for population structure in case-control genome-wide association studies, the status being analyzed as a quantitative phenotype. Chen *et al.* proved that this method is inappropriate and proposed a score test for the mixed logistic regression (MLR). However this test does not allow an estimation of the variants’ effects.

**Results:** We propose two computationally efficient methods to estimate the variants’ effects. Their properties are evaluated on two simulations sets, and compared with other methods (MLM, logistic regression). MLR performs the best in all circumstances. The variants’ effects are well evaluated by our methods, with a moderate bias when the effect sizes are large. Additionally, we propose a stratified QQ-plot, enhancing the diagnosis of *p*-values inflation or deflation, when population strata are not clearly identified in the sample.

**Availability:** All methods are implemented in the R package **milorGWAS** available at https://github.com/genostats/milorGWAS.

**Supplementary information:** Supplementary data are available at *Bioinformatics* online.

## 1 Introduction

Population stratification has long been known to be at the origin of spurious associations in genetic association studies (Lander and Schork, 1994): if the frequency of the phenotype of interest varies across the population strata, it will be associated to any allele the frequency of which varies accordingly. An early and elegant solution to this issue has been the use of family data, notably in the Transmission Disequilibrium Test (TDT) (Spielman *et al.*, 1993) and in the Family Based Association Test (FBAT) (Rabinowitz and Laird, 2000). However, these methods imposed the ascertainment and genotyping of affected individuals’ relatives, impairing their practical feasibility. The advent of Genome-Wide Association Studies (GWAS), demanding increasingly large samples to detect weaker and weaker effects, made the problem even more accurate. Methods adapted to large scale population studies have thus been proposed, among which the genomic control (Devlin and Roeder, 1999), which uses the empirical distribution of the genome-wide chi-square statistics to correct the statistic inflation attributable to population structure, and Structured Association methods, which first infer population strata from genome-wide data, then test for association conditional to the strata (Pritchard *et al.*, 2000b,a).

A major breakthrough was achieved in 2006 with EIGENSTRAT (Price *et al.*, 2006), also known as Principal Component Regression (PCR), a conceptually simple but extremely efficient method that consists in incorporating the top Principal Components (PC) of the genotype data in a linear model. It was then natural to use mixed models, as they can be interpreted as a generalization of PCR which incorporates all PCs in the model with random effects (Zhang and Pan, 2015; Dandine-Roulland and Perdry, 2015). Incorporating a few PCs with fixed effects as well in the model might still prove useful to correct statistic inflation at SNPs with large allelic frequency variations across strata (Price *et al.*, 2010; Zhang and Pan, 2015). Fast approximate (Aulchenko *et al.*, 2007) or exact (Lippert *et al.*, 2011) methods for genome-wide analysis of quantitative traits with mixed linear models (MLM) were soon made available. As fitting a mixed logistic regression (MLR) model remained computationally heavy, in case-control studies the status was coded as quantitative trait (0 or 1), and analyzed as such.

However, Chen *et al.* proved that when disease prevalence was heterogeneous between populations strata, while the overall distribution of *p*-values is well corrected by this method, it leads to conservatives *p*-values for some SNPs and to anti-conservative *p*-values for others. This behavior was made evident by the mean of quantile-quantile plots (QQ-plots) in which SNPs were categorized according to their allele frequencies in the different population strata. Chen *et al.* (2016) proposed a score test for the MLR, which is feasible in GWAS, and showed that the *p*-values obtained with this test were well distributed in all SNP categories.

While the score test has a reduced computational burden, its drawback is the absence of an estimation of the variants’ effects. When only the effects of genome-wide significant SNPs are needed, an obvious solution is to fit MLR models including each of these SNPs. When the SNPs’ effects are needed for the whole genome, e.g. for meta-analysis purposes, it is desirable to have a computationally efficient method to estimate these effects in mixed logistic regression. We propose two such methods in this paper.

One method is based on a first order approximation of the MLR, which leads to an approximation of the SNPs effect; the resulting Wald test is identical to the score test of Chen *et al.* (2016). A similar formula appeared without proof in Zhou *et al.* (2018). The other method, which bears similarities with the methods of Aulchenko *et al.* (2007), consists in first estimating individual scores in a mixed logistic regression model, and then incorporating these effects as an offset in a (non-mixed) logistic regression model.

We evaluate the capacity of logistic regression, MLM, and MLR (using either the score test or the methods mentioned above) to properly take into account population structure associated to heterogeneous prevalence. While Chen *et al.* were interested in geographically distinct populations, part of our work focuses on the context of two geographically very close populations in West Africa, using real genotype data from a recent GWAS (Milet *et al.*, 2019). We also use data simulated with a coalescent model (Hudson, 2002), reproducing the simulations presented in Chen *et al.* (2016). We use similar simulations to evaluate the ability of the PQL and of our two methods to properly evaluate SNPs’ effects.

Additionally, we propose to generalize the categorization of SNPs in QQ-plots presented in Chen *et al.* (2016) to variables other than the population strata, including continuous variables as for example the first PC. The interest of this generalization is demonstrated on the same simulations.

All methods are implemented in the R package **milorGWAS** (for **mi**xed **lo**gistic **r**egression in **GWAS**), freely available on github.

## 2 Methods

### 2.1 Fast methods for mixed logistic regression

The mixed logistic regression model (or logistic mixed model) considered is

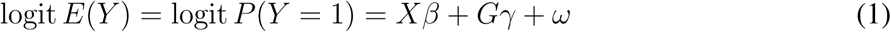

where *Y* is an *n*-dimensional vector of zeroes and ones (logit *E*(*Y*) is the vector of components logit *E*(*Y*_*i*_)), *X* is a *n* × *p* matrix of covariates (including a column of ones for the intercept), *G* is a vector of genotypes (ususally coded 0, 1 or 2), and *ω* is a random vector following a multivariate normal distribution *MV N* (0, *τ K*).

The likelihood of this model involves an integral over the random vector *ω*. There is no closed form for this integral, and numerical integration schemes are computationally intensive in high dimension. A classical approximate solution is the Penalized Quasi-Likelihood (PQL) algorithm, which is a sequence of approximations of the MLR model by linear mixed models. Even the PQL is too computationally intensive for estimating the effect *γ* of all SNPs in a GWAS.

We describe below two approximate methods for estimating *γ*. The first is based on an approximation of the maximum likelihood estimate in the PQL. The second consists in first estimating predicted linear scores 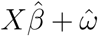 in the MLR model (1) without the term *Gγ*, and then incorporating these linear scores as an offset in a (non-mixed) logistic regression model.

#### 2.1.1 Approximate Maximum Likelihood Estimate (AMLE)

We outline here the general principle on which the formula presented in supplementary material can be derived. This principle could be applied to any statistical model. Let *ℓ*(*κ, γ*) be a log-likelihood, in which *κ* is a nuisance parameter and *γ* is the parameter of interest. The null hypothesis to be tested is *H*_0_ : *γ* = 0. Denote 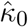 the Maximum Likelihood Estimator (MLE) of *κ* under the null hypothesis:

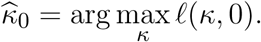

The score statistic to test for *H*_0_ is the first derivative in *γ* at the point 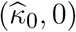 :

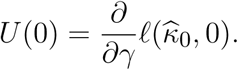

The null hypothesis can be tested using *T* = *U*(0)^2^*/* var(*U*(0)), which asymptotically follows a *χ*^2^ distribution. A second order approximation, with 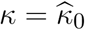 fixed, gives

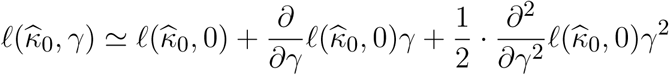

which maximizes in

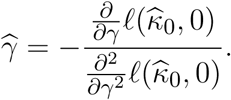

This estimator of 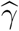 is not the MLE of *γ*, but when the true value of *γ* is small enough, both estimators are close.

In the context of GWAS, *κ* is the vector of random term variance and covariates effects, while *γ* is the effect of the SNP to be tested. This estimator shares with the score test the advantage that 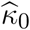 has to be estimated only once; the partial derivatives in *γ* are usually easy to compute, allowing a fast testing and estimating procedure.

However, as mentioned above, in the case of the MLR, the likelihood can’t be computed efficiently. We use the PQL to estimate the nuisance parameter *κ* = (*β, τ*), and the log-likelihood of the last linear approximation used in the PQL to estimate *γ*. The variance of the resulting 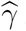 is estimated in this linear approximation; the resulting Wald test is identical to the score test of Chen *et al.* (2016). All details are given in the Supplementary Materials.

#### 2.1.2 Offset

The proposed method consists in estimating a vector of individual effects, including both the random components and the covariates in *X*, which is then incorporated in a logistic regression as an offset:

- First estimate 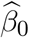 and 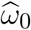 under the hypothesis *γ* = 0, in the MLR model

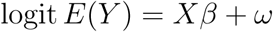

with *ω* as in (1).
- Then, for each vector of genotypes *G*, fit a linear model for

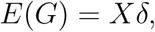

let 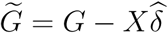 be the residuals of *G*, and estimate *γ* in the fixed-effects logistic regression

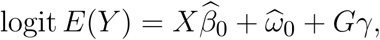

in which the vector 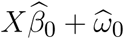 is an offset (that is, is held constant).

The motivation of this heuristic is that a similar two-steps method applied to a linear model *E*(*Y*) = *Xβ* + *Gγ* would give the same estimator 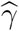 than the classical regression (cf Supplementary materials for the details).

### 2.2 Stratified QQ-plot

One of the contributions of Chen *et al.* was to show that a QQ-plot of log *p*-values was not sufficient to diagnose an incorrect test procedure, and to propose a “stratified QQ-plot” in which different categories of SNPs are represented separately. This allowed to see that in some of these categories, the test statistics are either inflated or deflated, while the overall distribution of *p*-values was correct. Here is how their categories were defined.

Chen *et al.* consider a population with two strata indexed by *i* = 0 or 1. The strata are assumed to be panmictic, so that expected variance of a SNP genotype *G* in stratum *i* is var_*i*_(*G*) = 2*p*_*i*_*q*_*i*_, *p*_*i*_ and *q*_*i*_ being the SNP allele frequencies. Each SNP is categorized according to the variance ratio *r*(*G*) = var_1_(*G*)*/* var_0_(*G*) between the two strata as follows (Chen *et al.* use a threshold *th* = 0.8):

- The SNPs with *r*(*G*) < *th* are category 1,
- the SNPs with *th* ≤ *r*(*G*) ≤ 1*/th* are category 2,
- the SNPs with 1*/th* < *r*(*G*) are category 3.

We propose to extend the method to stratify QQ-plots according to any covariate *Z*. If *G* ∈ {0, 1, 2}^*n*^ is the vector of genotypes, *Z* a vector with components in the range [0, 1], and ***1*** denotes a vector of ones, we let

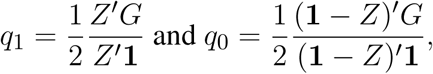

and we defined SNP categories as above (with *p*_*i*_ = 1 − *q*_*i*_). If *Z* is the indicator variable of the strata, *q*_1_ and *q*_0_ are the allelic frequencies in the two strata, and the categories will be identical to those of Chen *et al.*. The point of this extension is that when the relevant sub-strata are unknown, one could use one of the top genomic PCs instead (after rescaling them to [0, 1]).

### 2.3 Simulation studies: type I error in presence of population structure

We performed two sets of simulations, based on the simulations performed in Chen *et al.* (2016), to assess the efficiency of the different methods to correct for population stratification.

A crucial point of these simulations is the presence of two strata (or two cohorts) with different disease prevalence, and of related individuals. Simulations were performed under the null hypothesis of no genetic association (*γ* = 0 in equation (1)) and were analyzed with

- a logistic regression model
- a mixed linear model
- a mixed logistic regression model, using Chen *et al.* score test (identical to AMLE Wald test)
- a mixed logistic regression model, using the offset method

All analyses were repeated with the top ten PCs included as fixed effects in the model. We assessed the capacity of each test procedure to control type I error rates using Chen’s stratified QQ-plot. Moreover, to gauge the interest of our extension, we compared Chen’s QQ-plot to the stratified QQ-plot obtained using the first PC instead of the cohort indicator.

#### 2.3.1 Simulations based on South Benin data

We used genotype data from a GWAS on mild malaria susceptibility performed on two cohorts in South Benin (Milet *et al.*, 2019). The participants were ascertained in two sites distant of 20 km from each other, in three different health centers for the first cohort and two for the second one. After quality control (QC), the genotypes of 800 individuals were available, 525 in the first cohort, and 275 in the second one. The genotyping was performed with an Illumina HumanOmni5 chips (1,847,505 SNPs after QC and filtering out SNPs with minor allele frequency -MAF-less than 5%). This genetic sample presents both population structure and cryptic relatedness. Self-reported ethnic composition differed between the two cohorts, and principal component analysis confirmed the presence of population structure. A sub-structure related to the health center in which the participant was ascertained was also apparent. Moreover, substantial relatedness was observed in the sample, with levels of relationship corresponding to half-sibs, uncle-nephew or even 3/4 siblings for some pairs (estimated kinship coefficient *ϕ* from 0.10 to 0.16).

We simulated a binary phenotype with a difference of disease prevalence between the two cohorts, and a random effect modeling both population stratification and relatedness. Specifically, the probability on an individual *i* of being a case was calculated as:

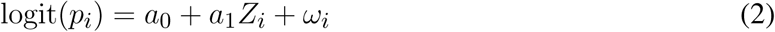

where *Z* is an indicator variable for belonging to the second cohort, and *ω*_*i*_ an individual random effect. The coefficients *a*_0_ and *a*_1_ were defined as *a*_0_ = logit(0.05) and *a*_1_ = logit(0.30) − logit(0.05), so as to obtain a prevalence of 0.05 in the first cohort and of 0.30 in the second one (not taking into account the presence of random effects). The vector of random effects was simulated following a multivariate normal distribution:

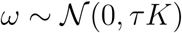

where *K* is the Genomic Relationship Matrix (GRM) calculated from all the SNPs. We set *τ* = 1.

#### 2.3.2 Simulations based on a coalescent model

We also performed coalescent simulations, reproducing closely the simulations described in Chen *et al.* (2016), to obtain genotypes for a large cohort of 10 000 individuals, with both population structure and relatedness, using the `ms` software (Hudson, 2002). This procedure based on a stepping stone model with symmetric migration between adjacent cells of the grid is commonly used to simulate a population with a spatially continuous population structure (Mathieson and McVean, 2012; Bradburd *et al.*, 2016). We use a 20 × 20 grid, in which the migration rate between adjacent cells was set to 10; this parameter produces a Wright’s fixation index *F*_*st*_ < 0.01 when dividing the simulated grid into two equal sub-populations, a level comparable to what is observed within Europe (Mathieson and McVean, 2012). We simulated a total of 10 millions independent SNPs. After filtering out SNPs with a MAF lower than 5%, 2 840 903 SNPs were available. The full command line arguments for `ms` are included in Supplementary Materials.

To obtain related individuals, we first simulated the genotypes for 8 000 founders (20 on each of the 400 cells). We then sampled 10 pairs of individuals in each cell, forming 4 000 couples, and simulated two offsprings by gene dropping. Thus we obtained 16 000 individuals (founders and offspring) from which 10 000 individuals were randomly selected to obtain the cohort.

The phenotype was simulated as before using equation (2), where *Z*_*i*_ was set to one when individuals were sampled in top left 10 × 10 grid, corresponding to a strata with a higher risk. The values of *a*_0_, *a*_1_ and *τ* were set as before; in this simulation set, *K* = 2Φ were Φ is the matrix of kinship coefficients (entries are 0.5 for first order relatives, 1 on the diagonal, 0 elsewhere). Data analyses were subsequently performed using a GRM calculed from 100 000 random SNPs.

### 2.4 Simulations studies: estimation of the SNPs’ effects

Simulations including a SNP effect were performed using the real data set described in 2.3.1, using this time

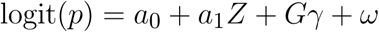

with *G* a genotype picked at random in the data, and *Z* and *ω* as described above. We considered four different scenarios:

A. Moderate cohort effect (respective prevalence *p*_0_ = 0.10 and *p*_1_ = 0.20) and moderate random effect (*τ* = 0.3).
B. Moderate cohort effect (respective prevalence *p*_0_ = 0.10 and *p*_1_ = 0.20) and large random effect (*τ* = 1).
C. Large cohort effect (respective prevalence *p*_0_ = 0.05 and *p*_1_ = 0.30) and moderate random effect (*τ* = 0.3).

The coefficients *a*_0_ and *a*_1_ are computed as *a*_0_ = logit(*p*_0_) and *a*_1_ = logit(*p*_1_) − logit(*p*_0_); *G* is centered to ensure that the expected prevalence is as prescribed. For each scenario, we considered SNPs with MAF in intervals (0.05; 0.10], (0.20; 0.25] and (0.45; 0.50], and SNP effect *γ* = log(1.5) and *γ* = log(2) (corresponding to *OR* = 1.5 and 2). One hundred replicates were performed for each condition, and analyzed with the PQL, the Offset and the AMLE, including the top 10 PCs.

## 3 Results

### 3.1 Type I error in presence of population structure

Figure 1 displays the stratified QQ-plots of the logistic regression, the mixed logistic regression (Chen’s score test, identical to AMLE Wald test) and the mixed linear model obtained on the genotype data from South Benin with a simulated phenotype, with or without the top 10 PCs. The 1 847 505 SNPs are splitted in three categories as described above (using a threshold *th* = 0.8). Category 1 contained 11.4% of the SNPs, category 2 and 3, respectively 77.6% and 11.0% of the SNPs.

**Fig. 1.**
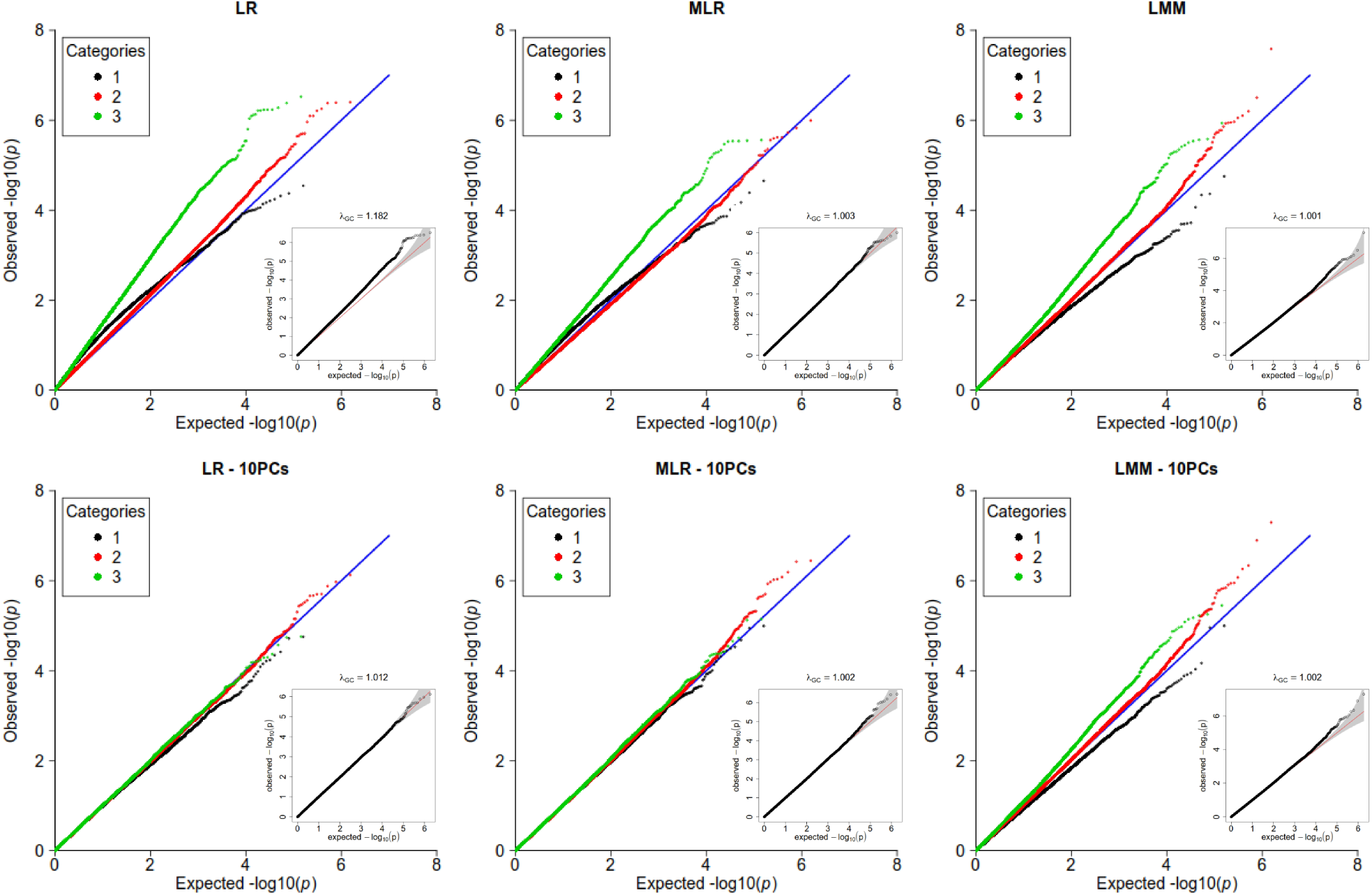
Stratified quantile-quantile plots for logistic regression (LR), mixed logistic regression (MLR) using Chen’s score test (or AMLE), and mixed linear model (MLM) on the data simulated based on South Benin data. In each panel, a non-stratified QQ-plot is embedded. On the second row, 10 PCs were included as covariates. SNP categories are determined as in Chen et al. (2016), based on the allele frequencies in the strata.

When no PCs are included (first row of the figure), statistic inflation is observed for the logistic regression (*λ* = 1.182). Based on the non-stratified QQ-plot, both MLM and MLR appear to adequately correct for population structure; however the stratified QQ-plot shows that this is not the case for SNPs in categories 1 and 3. In particular, for MLM, there is a statistic inflation for SNPs in category 3, and a deflation for SNPs in category 1.

When 10 PCs are included in the models (second row of the figure), this difference of behavior between SNPs categories persists for MLM. However, both logistic regression and MLR show an adequate correction for all categories of SNPs.

Similar comparisons were performed using the coalescent simulations of 10 000 individuals on a 20 × 20 grid, including first order relatives (Supplementary Figure 1). The 2840 903 independent SNPs are at 23, 7% in category 1, 58.8% in category 2 and 17.5% in category 3. The same patterns of inflation and deflation of the test statistic are retrieved for most cases; the only notable difference is that, in this case, the logistic regression including 10 PCs does not correct completely the statistics inflation.

Figure 2 shows the QQ-plots for the Offset method with the top 10 PCs included in the model, on the two simulations sets. While it adequately corrects for population structure in the data from the South Benin, it is too conservative in the case of the simulated cohort. simulations

**Fig. 2.**
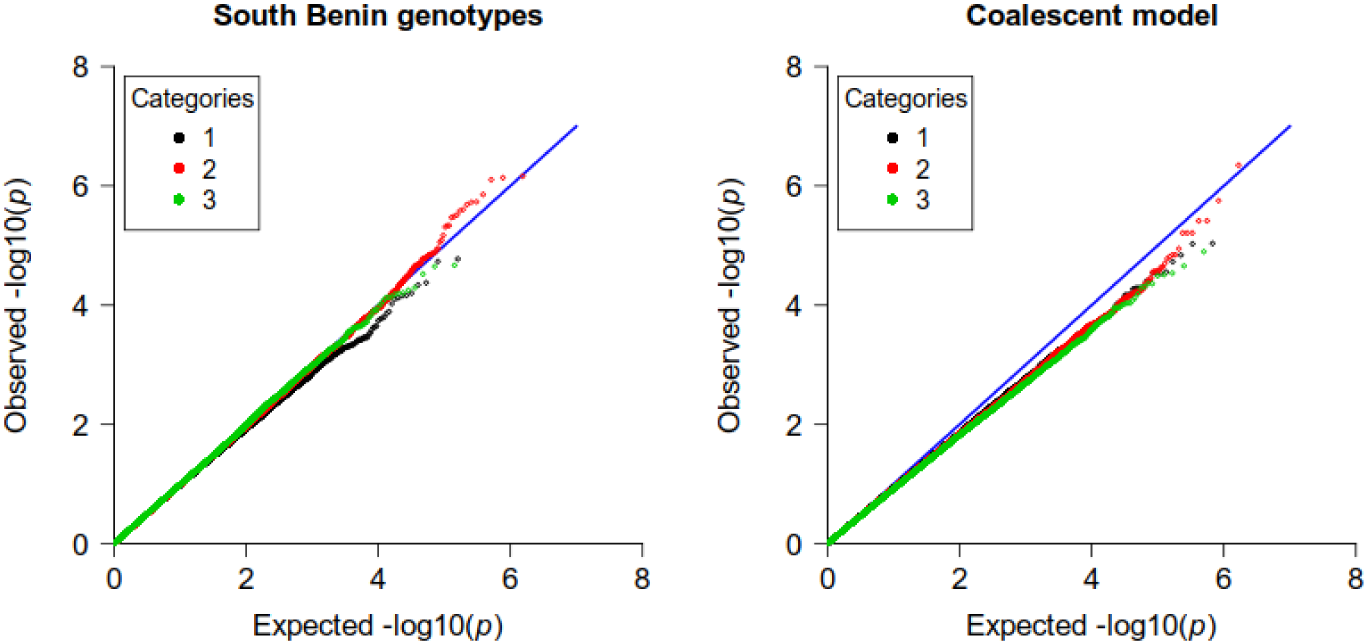
Stratified quantile-quantile plots for the Offset method, for simulations based on South Benin data (left) and on the coalescent model (right). SNP categories are determined as in Chen et al. (2016), based on the allele frequencies in the strata.

### 3.2 Extension of the stratified QQ-plot

Figure 3 compares the stratified QQ-plots obtained using either strata information (as in figures 1 and 2) or the first PC, for two of the analyses already considered in figure 1, the MLM and the MLR, both including 10 PCs as fixed effects. The same comparisons were performed for analyses on the simulated cohort (Supplementary Figure 2). While there are small differences between the QQ-plots, we see that they allow similar diagnostics, that is, an incomplete correction of population structure for MLM analyses, and in contrast an adequate correction for MLR.

**Fig. 3.**
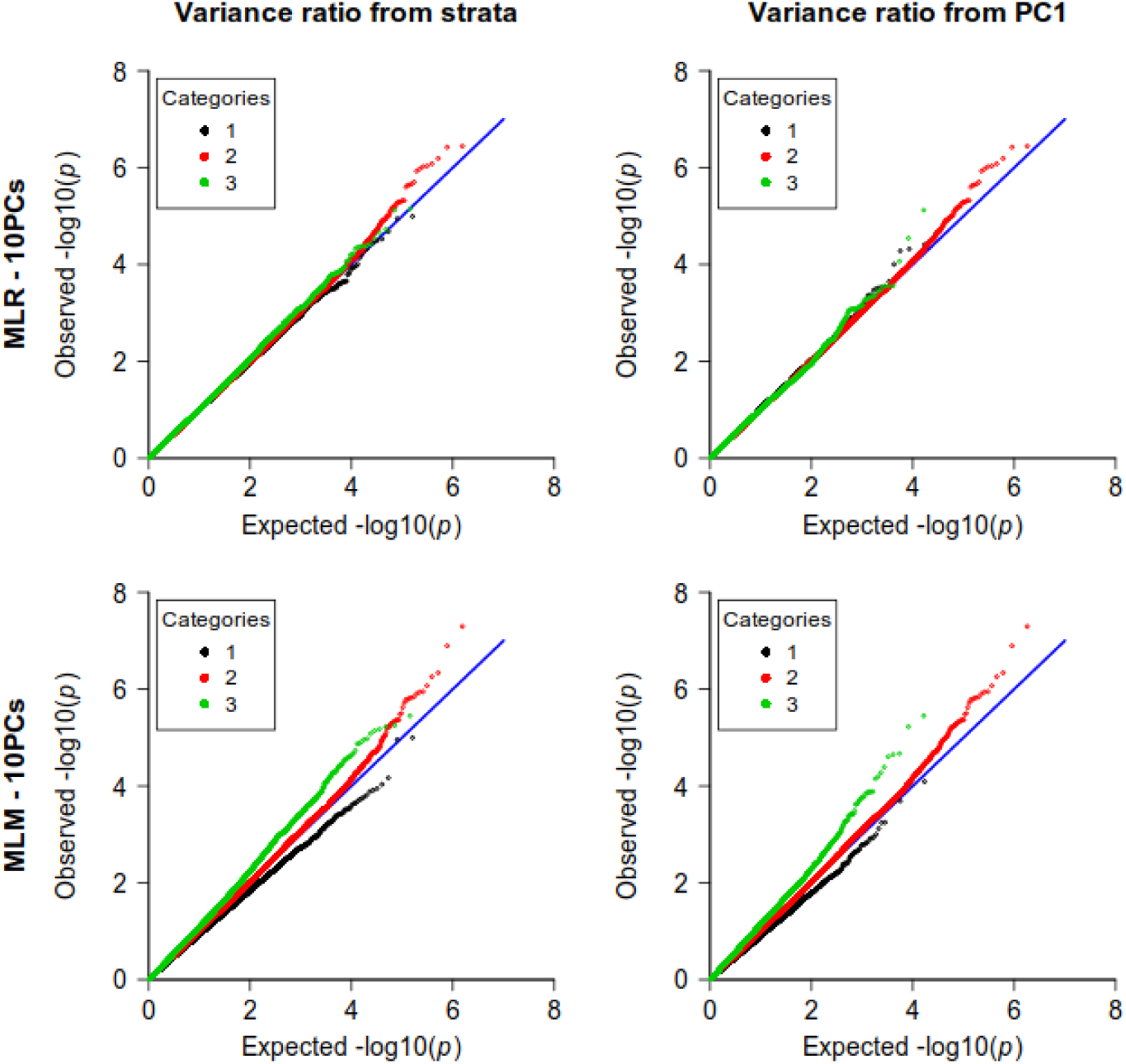
Stratified quantile-quantile plots obtained from the allele frequencies in the two strata (left) and from the first PC coordinates (right) for simulations based on South Benin data.

### 3.3 Estimation of the SNPs’ effects

Figure 4 shows the bias 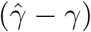 obtained for two different values of *γ*, in three scenarios A, B and C corresponding to different magnitude of cohort and random effects. Three MAF bins were considered (from 0.05 to 0.10, from 0.20 to 0.25 and from 0.45 to 0.50).

**Fig. 4.**
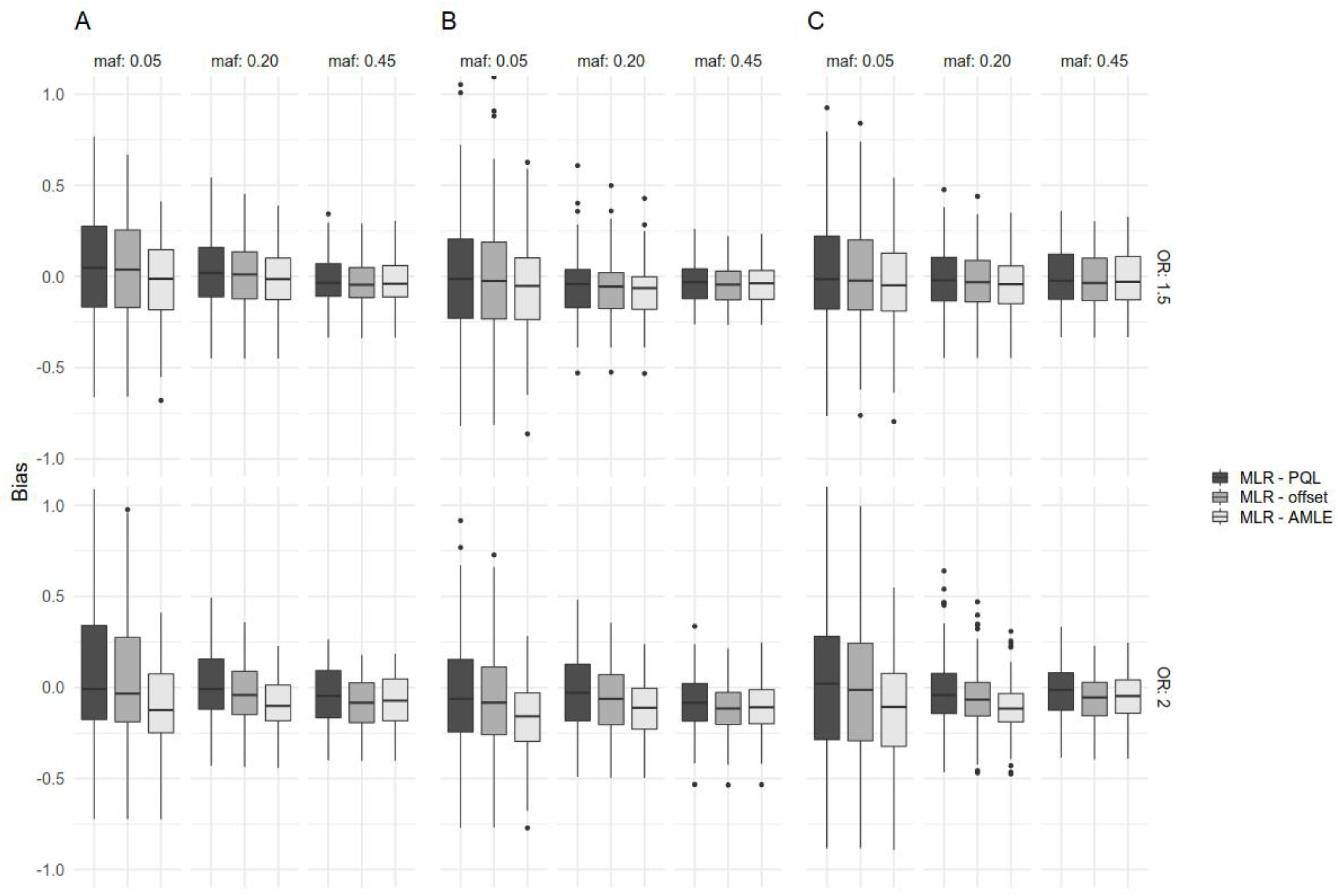
Bias of the SNP effect estimates 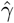 for two values of *γ* (*γ* = log(1.5) and *γ* = log(2)), in three scenarios (A: moderate cohort and random effect; B: moderate cohort effect, large random effect; C: large cohort effect, moderate random effect), and three MAF bins (0.05 to 0.10, 0.20 to 0.25, and 0.45 to 0.50).

In all situations, the PQL displays no bias, or a very small bias (for example in scenario B, corresponding to large random effects). The two proposed methods tend to have a negative bias. For data simulated with a moderate SNP effect *γ* = 0.4 (corresponding to *OR* = 1.5), they both have small negative biases (at most −0.08 corresponding to estimated *OR* = 1.4), independently of the importance of population structure (scenarios A, B and C).

For larger SNPs effects (*γ* = 0.7, corresponding to *OR* = 2), the bias increases, with AMLE having the larger bias, attaining in some situations −0.1 (corresponding to an estimated *OR* = 1.8). The bias is slightly more important for large random effects (scenario B).

## 4 Discussion

Our first result is a reproduction of the observation made by Chen *et al.* that is, in the presence of heterogeneity of disease prevalence between population strata, the mixed linear model (MLM) is inappropriate to analyse binary traits, leading to conservative *p*-values for some SNPs, and to anti-conservative *p*-values for others, depending on the ratio of expected genotype variance in the two strata. The motivation of Chen *et al.* was a genome-wide association study of asthma including individuals from different Caribbean and Latin America backgrounds, with in particular *ca.* 15% of individuals from Puerto Rico, in which the prevalence of asthma was much higher (25.6%) than in other populations (from 3.9 to 9.6%). We retrieved similar results in an analysis of a simulated phenotype with large differences of prevalence among strata, based on genotype data from two geographically close cohorts (*ca.* 20 km apart) from South Benin (Milet *et al.*, 2019), but with different self-reported ethnicities. Heterogeneity of prevalence may result from environmental factors (e.g. lifestyle, nutritional behavior, etc), and could occur frequently in association studies, thus making the analysis of binary traits with the MLM incorrect, in particular in populations with a high genetic diversity.

A similar result was retrieved with data simulated with a coalescence model on a square grid, the “high risk strata” consisting in the top left quarter of the grid; these simulations included many first order related individuals and a random effect based on the kinship matrix. The mixed logistic regression (MLR) including the top PCs as fixed effects is the only method to completely correct for population structure in both simulations. A classical logistic regression including the top PCs can however be worth considering, as this conceptually simpler method was efficient enough for the South Benin data, in which the level of relatedness, though high, is lower than in the simulated data.

It is worth to note that in both simulations sets, it was necessary to include the top PCs as fixed effects in the MLR to obtain correct type I error. The interest of including top PCs alongside with the random components as been noted before (Price *et al.*, 2010; Zhang and Pan, 2015).

The diagnosis of correctness of type I error cannot be based on the sole QQ-plot of *p*-values, as the behavior of the test differs in SNP categories defined from the allelic frequencies in the two strata, as mentioned above. When these strata are clearly identified in the study, Chen *et al.* introduced a QQ-plot stratified on SNP categories based on the allele frequencies in the strata. We propose an extension of this method that can be used when population information is not available, using the first PC as a proxy (or any continuous variable defined at population level, independently of the phenotype). Our simulations show that this method produces QQ-plot similar to the QQ-plot obtained with full knowledge of the two strata, thus allowing to diagnose whether the population structure is adequately taken into account or not.

We proposed two fast methods to estimate SNPs’ effects in mixed logistic regression. Both methods are conceptually simple and could be applied to other models (e.g. survival analysis), although literal computations to derive the formulas in the first one, AMLE, can be tedious. The Wald test performed with the AMLE is equivalent to the score test of Chen *et al.*, 2016, and thus the conclusions drawn for the MLR applied regarding type I error. The second method, which we call the Offset method, had similar performances on the simulations based on the South Benin genotypes, but was slightly over-conservative in the presence of strong familial effects in the simulations based on the coalescent model. We performed simulations based on the South Benin data to compare these methods with the PQL, which is the standard method for non-linear mixed models (although it is based on a an approximation), with inclusion of the top PCs. The two methods are slightly biased downward, the bias of the Offset method being less important than the bias of the AMLE, while the PQL has virtually no bias. It is known that in presence of unaccounted heterogeneity, logistic regression effect estimates have negative biases (Gail *et al.*, 1984; Cramer, 2007; Ayis, 2009); our result hints that the heterogeneity between population strata is not completely taken into account by these methods. However, the bias is sensible only for large effects such as *OR* = 2, which makes its impact virtually negligible in GWAS.

All methods are available in an R package **milorGWAS** based on the R package **Gaston** (Dandine-Roulland and Perdry, 2018) for data manipulation. The R and C++ source code of **milorGWAS** is available on github at https://github.com/genostats/milorGWAS.

## Acknowledgements

We thank the Genotoul bioinformatics platform Toulouse Midi-Pyrenees (Bioinfo Genotoul) for providing computing and storage resources.

## References

Aulchenko, Y. S. et al. (2007). Genomewide rapid association using mixed model and regression: a fast and simple method for genomewide pedigree-based quantitative trait loci association analysis. Genetics, 177(1), 577–585.

Ayis, S. (2009). Quantifying the impact of unobserved heterogeneity on inference from the logistic model. Communications in Statistics—Theory and Methods, 38(13), 2164–2177.

Bradburd, G. S. et al. (2016). A spatial framework for understanding population structure and admixture. PLoS genetics, 12(1), e1005703.

Chen, Z. et al. (2016). Testing for association in case-control genome-wide association studies with shared controls. Statistical methods in medical research, 25(2), 954–967.

Cramer, J. S. (2007). Robustness of logit analysis: Unobserved heterogeneity and mis-specified disturbances. Oxford Bulletin of Economics and Statistics, 69(4), 545–555.

Dandine-Roulland, C. and Perdry, H. (2015). The use of the linear mixed model in human genetics. Human heredity, 80(4), 196–206.

Dandine-Roulland, C. and Perdry, H. (2018). Genome-wide data manipulation, association analysis and heritability estimates in R with Gaston 1.5. Hum Hered, 83.

Devlin, B. and Roeder, K. (1999). Genomic control for association studies. Biometrics, 55(4), 997–1004.

Gail, M. H. et al. (1984). Biased estimates of treatment effect in randomized experiments with nonlinear regressions and omitted covariates. Biometrika, 71(3), 431–444.

Hudson, R. R. (2002). Generating samples under a Wright–Fisher neutral model of genetic variation. Bioinformatics, 18(2), 337–338.

Lander, E. S. and Schork, N. J. (1994). Genetic dissection of complex traits. Science, 265(5181), 2037–2048.

Lippert, C. et al. (2011). FaST linear mixed models for genome-wide association studies. Nature methods, 8(10), 833.

Mathieson, I. and McVean, G. (2012). Differential confounding of rare and common variants in spatially structured populations. Nature genetics, 44(3), 243.

Milet, J. et al. (2019). First genome-wide association study of non-severe malaria in two birth cohorts in Benin. Human Genetics.

Price, A. L. et al. (2006). Principal components analysis corrects for stratification in genome-wide association studies. Nature genetics, 38(8), 904.

Price, A. L. et al. (2010). New approaches to population stratification in genome-wide association studies. Nature Reviews Genetics, 11(7), 459.

Pritchard, J. K. et al. (2000a). Association mapping in structured populations. The American Journal of Human Genetics, 67(1), 170–181.

Pritchard, J. K. et al. (2000b). Inference of population structure using multilocus genotype data. Genetics, 155(2), 945–959.

Rabinowitz, D. and Laird, N. (2000). A unified approach to adjusting association tests for population admixture with arbitrary pedigree structure and arbitrary missing marker information. Human heredity, 50(4), 211–223.

Spielman, R. S. et al. (1993). Transmission test for linkage disequilibrium: the insulin gene region and insulin-dependent diabetes mellitus (IDDM). American journal of human genetics, 52(3), 506.

Zhang, Y. and Pan, W. (2015). Principal component regression and linear mixed model in association analysis of structured samples: competitors or complements? Genet Epidemiol, 39(3), 149–155.

Zhou, W. et al. (2018). Efficiently controlling for case-control imbalance and sample relatedness in large-scale genetic association studies. Nature genetics, 50(9), 1335.

